# CAIM: Coverage-based Analysis for Identification of Microbiome

**DOI:** 10.1101/2024.04.25.591018

**Authors:** Daniel A. Acheampong, Piroon Jenjaroenpun, Thidathip Wongsurawat, Alongkorn Krulilung, Yotsawat Pomyen, Sangam Kandel, Pattapon Kunadirek, Natthaya Chuaypen, Kanthida Kusonmano, Intawat Nookaew

**Author notes:** Influenza Research Institute, Department of Pathobiological Sciences, School of Veterinary Medicine, University of Wisconsin-Madison, Madison, WI 53711, USA.

## Abstract

Accurate taxonomic profiling of microbial taxa in a metagenomic sample is vital to gain insights into microbial ecology. Recent advancements in sequencing technologies have contributed tremendously toward understanding these microbes at species resolution through a whole shotgun metagenomic (WMS) approach. In this study, we developed a new bioinformatics tool, CAIM, for accurate taxonomic classification and quantification within both long- and short-read metagenomic samples using an alignment-based method. CAIM depends on two different containment techniques to identify species in metagenomic samples using their genome coverage information to filter out false positives rather than the traditional approach of relative abundance. In addition, we propose a nucleotide-count based abundance estimation, which yield lesser root mean square error than the traditional read-count approach. We evaluated the performance of CAIM on 28 metagenomic mock communities and 2 synthetic datasets by comparing it with other top-performing tools. CAIM maintained a consitently good performance across datasets in identifying microbial taxa and in estimating relative abundances than other tools. CAIM was then applied to a real dataset sequenced on both Nanopore (with and without amplification) and Illumina sequencing platforms and found high similality of taxonomic profiles between the sequencing platforms. Lastly, CAIM was applied to fecal shotgun metagenomic datasets of 232 colorectal cancer patients and 229 controls obtained from 4 different countries and primary 44 liver cancer patients and 76 controls. The predictive performance of models using the genome-coverage cutoff was better than those using the relative-abundance cutoffs in discriminating colorectal cancer and primary liver cancer patients from healthy controls with a highly confident species markers.

**Key Points:**

- Metagenomic coverage is an important index to obtain highly accurate species identification by reducing false positives from whole shotgun metagenomic data.
- Comparative analyses of CAIM and other bioinformatics tools for species identification on many mock community whole shotgun metagenomic datasets generated by short-read and long-read sequencing and synthetic datasets were performed, showing that CAIM has a very good performance compared with the other tools.
- Using the metagenomic coverage approach through CAIM improves the predictive power of species biomarkers identified from in stool samples of colorectal cancer and primary liver datasets.

## INTRODUCTION

Advances in sequencing technologies have greatly contributed to the understanding of microbes found within diverse environments [1]. Various technologies have been used to sequence genetic material purified from microbial communities. Sequencing of metagenomic samples has efficiently aided researchers’ understanding of the role of these microbes in the microbiota, thus leading to a tremendous improvement in areas such as health care and agriculture yield [2, 3]. Metagenomic studies are often conducted in clinical and environmental samples to detect, identify, and enumerate the microorganisms present in a sample [4]. Whole shotgun metagenome sequencing (WMS), or the sequencing of all the genomic DNA in a microbiota sample, provides better resolution than 16S rRNA gene sequencing [5], and through the use of such techniques, analysis of complex environment samples has become possible using high-throughput sequencing technologies and bioinformatics tools.

Due to the complexities associated with metagenomic samples and the high volume of sequenced data they produce, systems involved must be fast and able to handle large amounts of data while accurately identifying species present within the samples. Several metagenomic classifier tools have been developed and benchmarked to aid in the identification of species and their abundance in the samples [6–8]; however, each method comes with its own challenges, including biases in the sequencing technologies, experimental errors, computational difficulties, and outdated reference databases [5, 6, 9]. False negatives and false positives are also common problems associated with most tools [7, 9–11]. Expansion in the number of genomes in reference databases has contributed to the identification of previously unidentified species and strains present in metagenomic samples, which would not be captured otherwise [12, 13]. Nevertheless, greater numbers of reference genomes in the database can increase the number of false positives associated with most tools in taxonomic classification [13].

Relative abundance level has been widely used as a cutoff criterion by most tools to filter out noises in metagenome data [7]; however, identifying the appropriate cutoff, which maximizes the accuracy in samples, is challenging due to the high diversity of microbial communities across different ecological systems. The lack of consensus on the relative-abundance cutoff in filtering out false positives poses a hindrance to accurately identifying the taxa present in microbiome samples, especially when dealing with complex microbiota [14]. For instance, in most staggered mock community datasets (ex. ZymoBIOMICS Microbial Community Standard II (Log Distribution) (Cat No. D6310), Zymo research, USA), the common relative-abundance cutoffs used by most existing tools (0.01 or 0.001) could overlook many species, as four out of the ten species have a low percentage of genomic DNA (<0.01). This is of concern for scarce but potentially important species in a community or low microbial biomass studies [15]. In addition, relative abundances are often estimated from the read counts mapped on the genome sequence of individual identified taxa; thus, genomes with fewer read counts are often excluded using the abundance-cutoff approach.

We developed an alignment-based metagenomic classification and quantification tool, <u>C</u>overage-based <u>A</u>nalysis for Identification of <u>M</u>icrobiome (CAIM), for the taxonomic classification of both short- and long-read WMS data. A further reduction in the reference genome datasets before alignment and the use of the genome coverage (the proportion of genome size covered by the aligned reads) in filtering out false positives could lead to high confidence in the microbial taxon identified as compared to the relative abundance. Moreover, the use of the sequencing depth of a microbial taxon to estimate the relative abundance provided a better estimate as compared to the read counts. We first evaluated the performance of CAIM using several mock community and synthetic datasets to compare it to the most advanced tools designed for metagenomic analysis of short reads (Metalign [14], KMCP [16], and StrainPro [17]), and both short and long reads (Kraken2 [18]). Limited number of previous studies have demonstrated consistent results when comparing short- and long-reads sequenced on distinct platforms, despite variations in mapping coverage and depth on WMS analysis [19, 20]. Therefore, we subsequently applied CAIM to evaluate the concordance between short- and long-read WMS data derived from the same aliquot of a multispecies purified fecal DNA samples. We further applied CAIM on real metagenomic samples collected from colorectal cancer (CRC) [21–23] and liver cancer [24] patients and healthy controls from four different countries to identify the predictive species biomarkers.

## METHODS

Methods are described in the Supplementary Methods.

## RESULTS

### Representative reference genomes

Reference genome collection and harmonization is a critical step to obtain comprehensive reference genomes with correct taxonomic labeling. Therefore, we collected high-quality genomes with taxonomic annotation from various databases, including 29,004 prokaryote genomes from the Genome Taxonomy Database [25], 1,635 fungi genomes from Joint Genome Institute [26], and 119 fungi genomes from the National Center for Biotechnology Information (NCBI) RefSeq (genomes) [27], and 10,628 viral genomes from the NCBI RefSeq database. We focused on species-level identification; therefore, after filtering for redundancies, 41,386 genomes were represented in the reference genome set (RRGS)—1,346 archaea genomes, 27,658 bacteria genomes, 1,754 fungi genomes, and 10,628 viral genomes.

### CAIM computational workflow for species classification and quantification of short and long reads WMS

CAIM workflow (Fig. 1) begins with the high-quality WMS data analysis of raw reads, quality control, and filtering contaminated host DNA reads to obtain high-quality reads for further steps.

**Figure 1.**
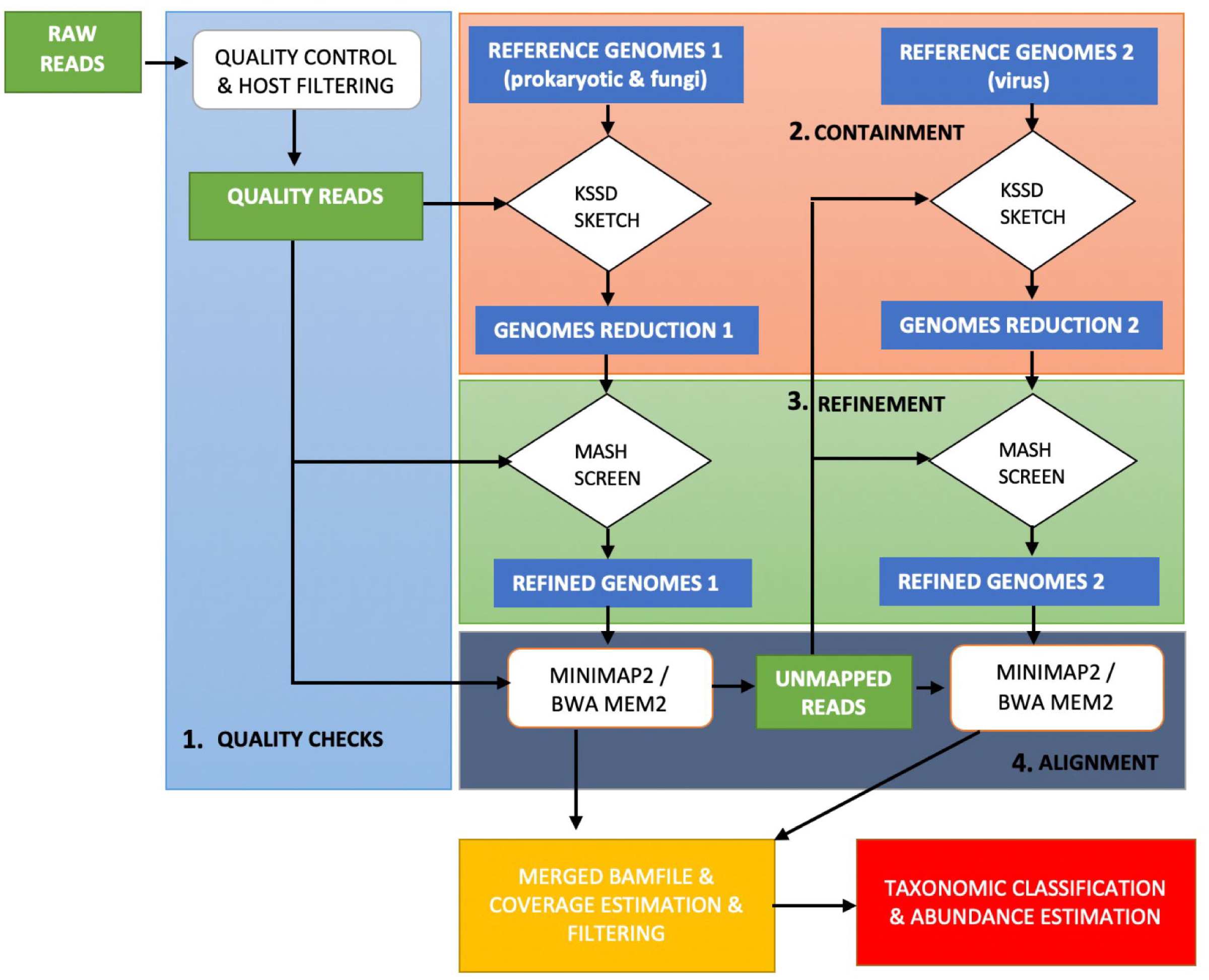
CAIM workflow: (1) High-quality metagenomic reads are screened against sketched reference genomes. The reference genomes are further reduced (2) and refined (3) which is later mapped against the metagenomic reads. (4) Unmapped reads are sketched and screen against Reference genomes to obtain refined genomes and later aligned with the unmapped reads. The aligned results are then merged for taxonomic classification. The light blue, red and green is for quality, containment and refinement steps respectively.

The second step, sample-dependent reference genome reduction, is important for reducing false positives and speeding up the read alignment. This process eliminates reference genomes of non-resemblance species in the considered sample from high-quality reads using containment analysis of sketching technique tools, k-mer substring space decomposition (Kssd) [28], and Mash screen [29], as k-mer information depends on genome size [30]. We split RRGS into two subsets—RRGS1 (prokaryotic and fungi genomes) and RRGS2 (viral genomes). We applied two steps of reference genome reduction by first sketching and filtering the reference genome based on containment scores calculated by Kssd. Kssd is faster than Mash but less efficient in sketching small-size genomes, resulting in inconsistent detection of species [26]. We then applied a second round of refinement based on containment scores calculated by Mash. This two-step approach gives a better consistency of reference genome reduction. We refined RRGS1 first, then repeated the process with RRGS2 using the unmapped reads, considering that the viral content in metagenomic samples is typically much smaller than that of prokaryotes and fungi.

The third step is classification and quantification. The high-quality reads were aligned to the refined RRGS1 then the unmapped reads were aligned to RRGS2 using Minimap2 [31] and Bwa-mem2 [32] for long and short reads. Sequencing depth and genomic coverage play a crucial role in estimating the relative abundance and identification of organisms in the metagenomic sample; [33] therefore, we implemented functions to calculate genome coverage, sequencing depth, and nucleotide count relative abundance using samtools [34] and bedtools [35]. The genome coverage was used as the important cutoff for filtering out false positives in species identification. The traditional way to calculate relative abundance relies on read count, which ignores the length of alignments that could vary in different reads, especially in long reads. The nucleotide-count relative abundance was calculated using the number of nucleotides of reads aligned to the individual genome relative to the number of total nucleotides of reads aligned on the refined RRGSs. The CAIM pipeline is packaged in Nextflow [36].

### The nucleotide-count approach performed better than the read-count approach for estimating relative abundance

We compared the performance of our proposed nucleotide count approach with the traditional approach of read count on estimation of relative abundance using several mock community datasets of different kingdoms, relative abundance profiles, and sequencing platforms (Fig. 2A). Only the genomes of species present in each mock community dataset (the perfect reference genomes) were used for read alignments. We then calculated the relative abundances from the alignment results using either read count or the nucleotide count and compared the calculated relative abundance with known relative abundance using root mean square error (RMSE) (Fig. 2C). A lower RMSE is preferred, as it implies that a particular approach’s accuracy is good relative to the ground truth abundance. In most cases, the nucleotide count approach gave lower RMSE than the read count approach across multiple mock community datasets for both short reads (ATCC MSA-1000, ATCC MSA-1001, ATCC MSA-1002, ATCC MSA-1003, Gut-HiLo, Gut-Mix, MCA-G2, MCA-N1, MCA-S1, MCB-N1, and MCB-S1) and long reads (ATCC MSA-1003, ATCC MSA-1003P, ATCC MSA-1002, BBMock-12, GIS20, Zymo-EVEN, ZymoQ20, and ZymoR103). RMSE of the read count approach was slightly better in a few of the mock community datasets for short reads (BBMock-12, GIS20, MCB-G1, and ZymoIL) and long reads (Zymo-LOG and ZymoD6331). There were considerably high RMSE (>5) associated with some of the short-read (MCA-G2, MCB-S1, MCA-N1, MCB-N1) and long-read mock community datasets (ZymoR103, ATCC MSA-1002). Overall, these results indicated that the proposed nucleotide count approach performs better than the widely used read count approach.

**Figure 2.**
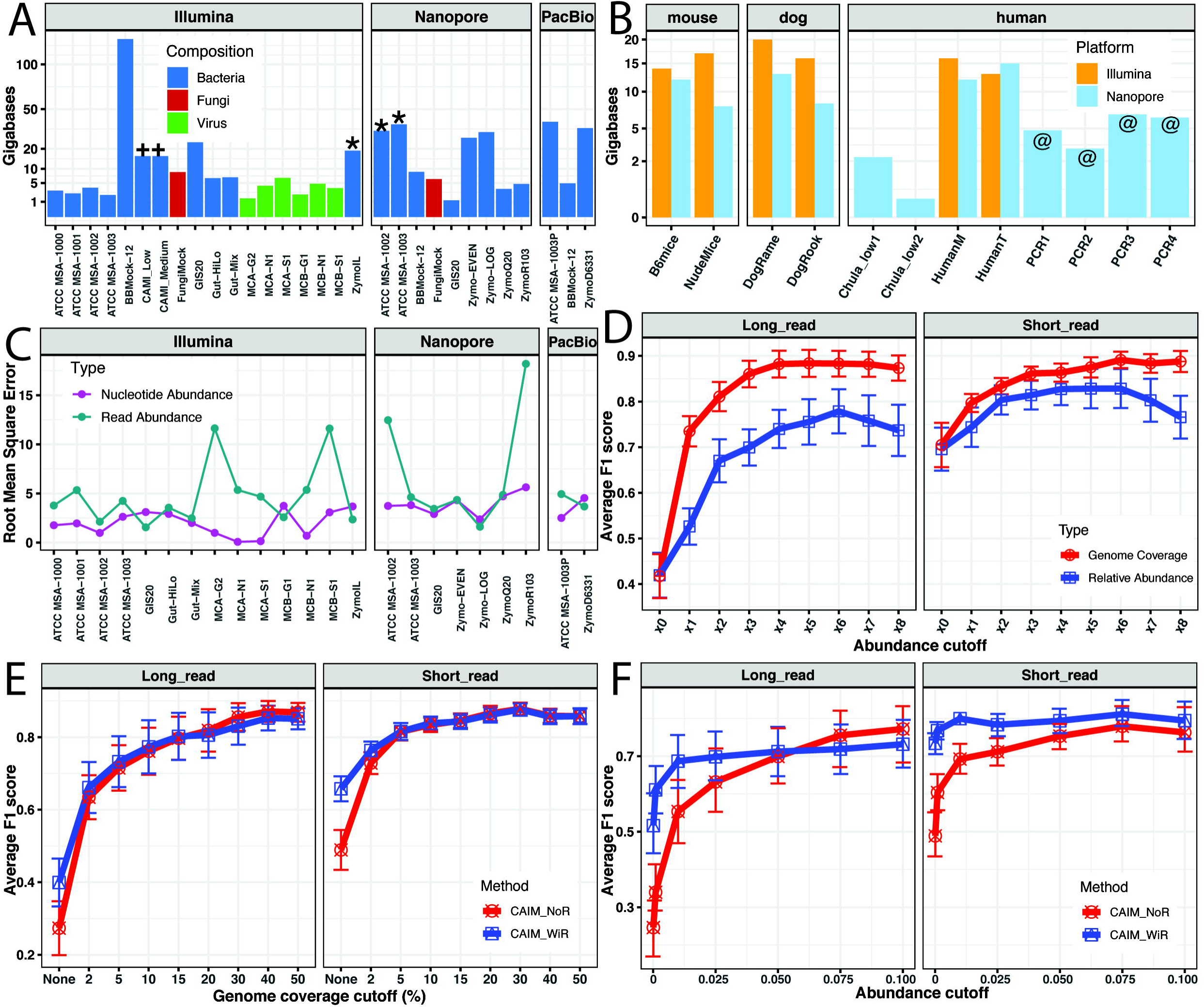
Bar plot showing the sizes of mock community (A) and real datasets (B) used in this study. Asterisks (**\***) represent mock community datasets sequenced in our lab and the plus signs (**+**) represent synthetic datasets. All the real datasets sequenced in our lab of native metagenome DNA amplified metagenome DNA indicated by the at sign (@). (C) Measure of root mean square error comparison between the nucleotide and read-abundance based on perfect reference genome scenario. (D) Average F1 scores based on the genome-coverage cutoff and relative-abundance cutoff for short reads and long reads on different subsampled mock community datasets. (E-F) The average F1 score for all the mock datasets at different genome-coverage and relative-abundance cutoffs for CAIM with and without the refinement step in Fig. 1. (x_O_, x_1_, x_2_, x_3_, x_4_, x_S_, x_6_, x_7_, x_8_) corresponds to abundance cutoff values of (0.0001, 0.001, 0.01, 0.025, 0.05, 0.075, 0.1, 0.5, 1) and genome coverage cutoff of (*none*, 2%, 5%, 10%, 15%, 20%, 30%, 40%, 50%) CAIM_NoR (CAIM without refinement), CAIM_WiR (CAIM with refinement)

### Genome-coverage cutoff controlled false positives better than the relative-abundance cutoff for taxonomic identification in WMS

Relative abundance has been used as the standard procedure to filter out false-positive species identifications from WMS data. We proposed using genome coverage in filtering out false positives within the CAIM pipeline. We compared the performance of species identification by genome coverage versus relative abundance (nucleotide count) using F1 score across the different mock community datasets at different sequencing depths of individual datasets (see supplementary table S1 for the mock communities included in this study) (Fig. 2D). The mean F1 score across the different selected genome-coverage cutoffs (0%-50%) showed superior performance compared to the selected relative-abundance cutoffs (0.0001%–1%) in both long- and short-read datasets, and the average F1 score using the genome-coverage cutoff values were higher than using the relative-abundance cutoff. For both long and short metagenomic reads, a genome-coverage cutoff value of 5% had a higher average F1 score than when we used any of the relative-abundance cutoffs (Fig. 2C). A genome-coverage cutoff of 15% resulted in the highest F1 score for the long read metagenomic samples, whereas a genome-coverage cutoff of 30% resulted in the highest F1 score for the short read metagenomic samples.

To assess the impact of the refinement step on reference genome reduction in our workflow (Fig. 1), we compared the average F1 score derived from either the relative abundance (nucleotide count) or genome-coverage cutoff on all the mock datasets (Fig. 2E and F). CAIM with the refinement step (CAIM_WiR) had a better average F1 score than CAIM without the refinement step (CAIM_NoR) across different cutoffs of relative abundance. Interestingly, the mean F1 score values between CAIM_WiR and CAIM_NoR for both long- and short-read metagenomic samples when the genome-coverage cutoff was applied, were higher, indicating the robustness of genome coverage in filtering out false positives. Therefore, the genome-coverage cutoff will be the default cutoff for CAIM.

### CAIM with specified genome coverage has better performance in species identification than other bioinformatic tools when analyzing mock community and synthetic datasets

We further evaluated the species-identification performance of our CAIM method on the mock community and the synthetic datasets and compared it with that of other top taxonomic identification tools used for WMS analysis, such as Metalign [14], KMCP [16], Kraken2 [18], and StrainPro [17]. We also considered the scenario where we vary the relative abundance (CAIM_abun). CAIM using the genome-coverage cutoff of 15 %, had a better performance of higher F1 score in identifying the true microbial taxa in all the datasets of short-reads and long-reads (Fig. 3A and B, repectively).CAIM_abun had a good performance like the other tools.

**Figure 3.**
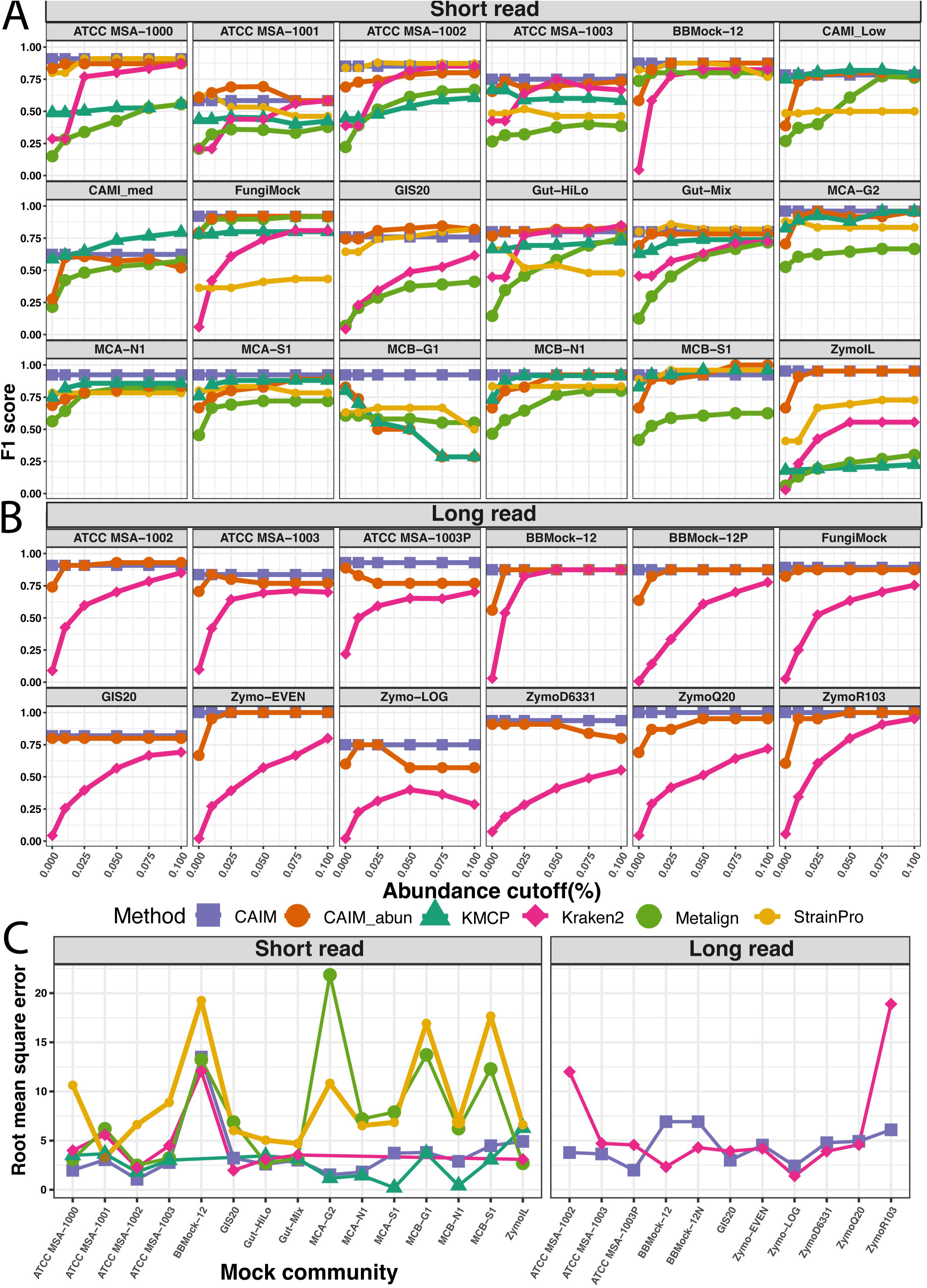
Comparison of different classifiers on short-(A) and long-read (B) metagenomic mock samples by estimating their F1 score at various relative-abundance cutoffs (%), while genome coverage cutoff of 15% was used for CAIM. (C) The root mean square error plot for the various taxonomic classifiers on the mock community datasets based on the higest F1 score from the previous panels.

We also compared the ability of CAIM to estimate the relative abundance in comparison with the other classifiers based on the highest F1 score of individul tools (Fig. 3C). The relative abundance estimated by CAIM using the nucleotide count approach performs equal to the theoretical abundance in most of the datasets. CAIM had the lowest RMSE using mock datasets in most scenarios compared to the relative abundance estimated by the other tool. There were a few cases where CAIM overestimated or underestimated the abundances. KMCP, Kraken2 performed equally well among themselves in most cases in estimating the abundances. Metalign, and StrainPro gave higher RMSE comparing to the other tools.

### High Concordance in species identification from WMS between Illumina and Nanopore sequencing platform

We applied CAIM to the real metagenomic samples and assessed concordance between short- and long-reads. We collected stool microbiome samples from humans, mice, and dogs and sequenced aliquots of each sample with Illumina and Nanopore sequencing platforms to create a dataset for the evaluation. By sequencing the same samples on the two different platforms, we investigated whether the species composition and their relative abundances from the two sequencing platforms are comparable, as well as the distinction of gut microbiome species across different hosts. The number of species identified from the gut microbiome of organisms of the same kind were similar, and the Spearman correlation between the relative abundance of common species identified within the same samples sequenced on different platforms (Fig. 4A) was high (rho ≥ 0.80).

**Figure 4.**
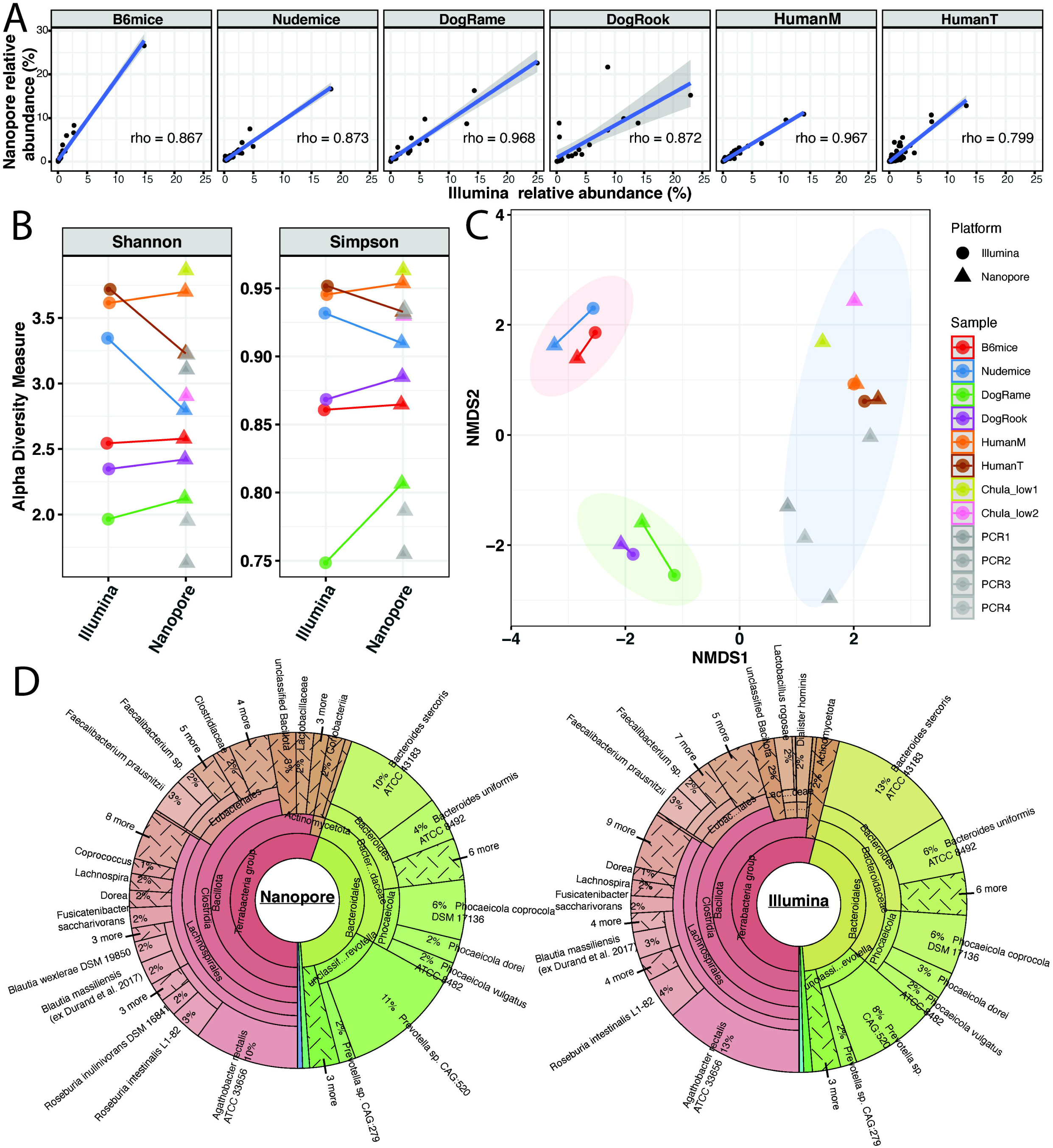
Comparison of species composition abundance in common from the gut microbiome of different organisms sequenced on both the Illumina and Nanopore sequencing platform as seen in scatter with Spearman’s rho correlation coefficient (A), Alpha diversity (B), and Beta diversity (C) plots. (D) Kronaplots of taxonomic compositions of sample HumanM showed a high simmilarlity between Illumina and Nanopore sequencing platform

We computed the Shannon and Simpson species evenness and degree of concentration of the two sequencing platforms, and the species richness and diversity of the Nanopore platform were slightly higher than the Illumina platform sample-wise (Fig. 4B). We compared the microbial diversity among samples of the same kind between the sequencing platform using the Bary-Curtis distance based non-metric multidimensional scaling (NDMS) and observed the samples clustered together more closely or separated by the gut microbiome of organisms of the same kind irrespective of the sequencing platforms and clustered together based on their kind except for the B6 and Nude mice which were slightly apart from each other (Fig. 4C). Using CAIM, we also investigated whether shallow (Chula_low2, Chula_low1) sequencing depth affected detected species composition or PCR amplification (PCR1,2,3,4) for human metagenomic samples. As expected, a higher number of species was identified for the sample with high sequencing depth as compared to the low sequencing depth as reported in the previous studies [37]. Nonetheless, beta diversity analysis of the community structure of human samples showed they were grouped together (Fig. 4C). The high simmilarlity of taxonomic profiles of the same aliquot derived from Nanpore and Illumina sequencing platform can be be observed as example of sample HumanM illustrated by kronaplots (Fig. 4D)

### Genome coverage improves the reliability and predictive power of CRC and primary liver cancers identification based on WMS

Several taxonomic classifiers and machine learning techniques have been applied to CRC metagenomic samples to identify microbiomes that help discriminate healthy controls from CRC patients [21, 23, 38]. Using three commonly used predictive modeling (random forest [RF], support vector machine [SVM], and least absolute shrinkage and selection operator [LASSO]) and three scenarios (within datasets, cross prediction, and leave one dataset out [LODO]), we investigated the predictive performance of using the genome-coverage cutoff approach (CAIM) and the relative-abundance cutoff approach (CAIM_abun) on datasets from four countries (used in previously publications [21, 36, 37]) to discriminate between healthy controls and CRC patients. For the within-datasets scenario, CAIM outperformed CAIM_abun in 10 out of the 12 cases (Fig. 5A) with an average AUC of 0.79 and 0.76, respectively. CAIM also outperformed CAIM_abun for the cross-prediction scenario in most of the cases for relative abundance (32 out of 36 cases) (Fig. 5A) and average model AUC (Fig. 5B,C). On average, the cross-prediction models using CAIM outperformed CAIM_abun among all the different models considered, apart from the SVM model prediction for GERMANY samples, which had the same average AUC value of 0.74. For the LODO scenario, CAIM had the best prediction AUC value (11 out of 12 cases) as compared to CAIM_abun among the three predictive models considered.

**Figure 5.**
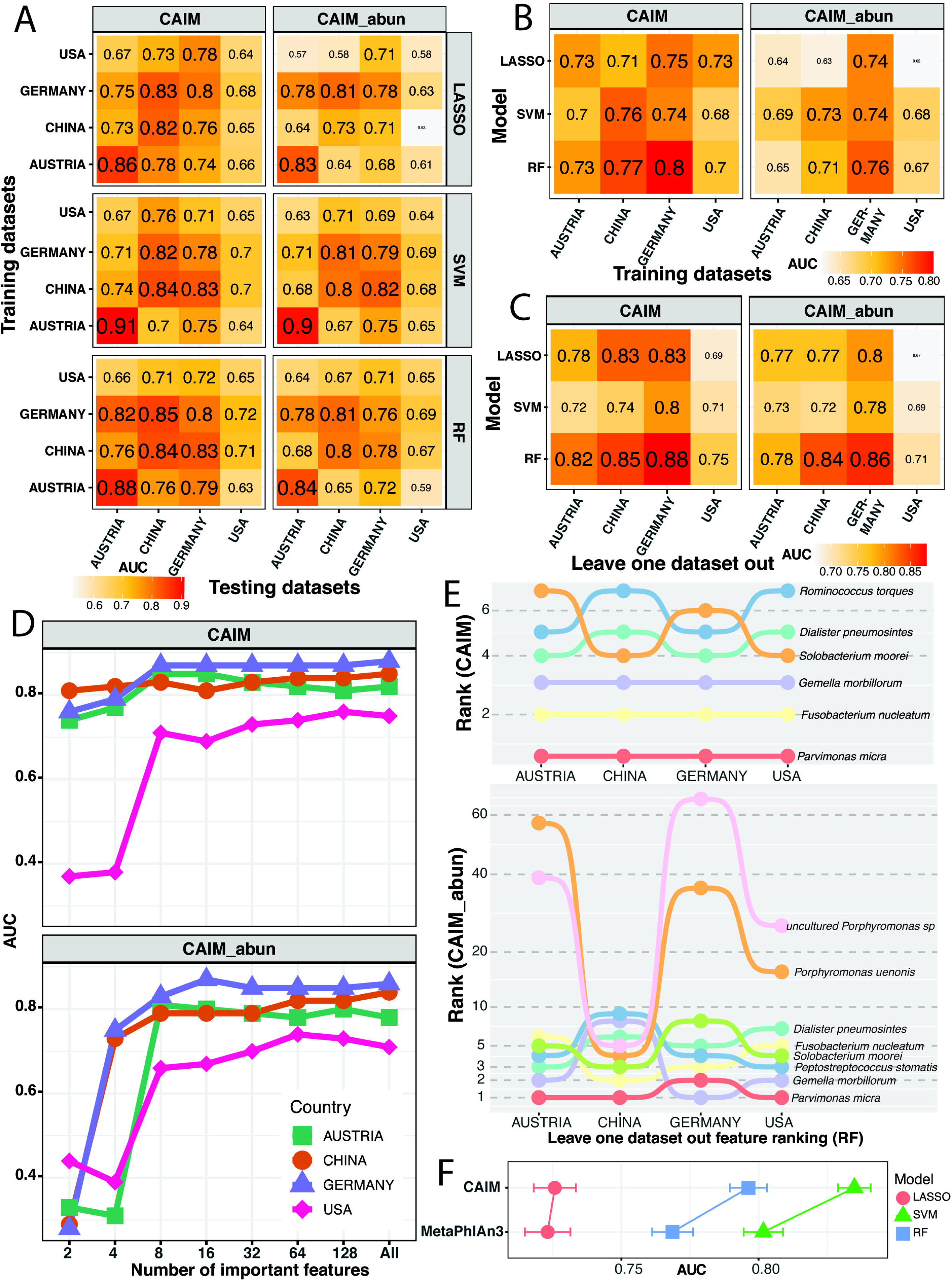
Predictive performance of random forest (RF), support vector machine (SVM), and least absolute shrinkage and selection operator (LASSO) predictive models using taxonomic profiles derived from CAIM and CAIM_abun relative abundances. (A) Cross-prediction models where we trained the model on the CRC datasets on the y-axis and tested on the x-axis datasets. The diagonal represents the within-dataset prediction where we trained the model on 80% of the datasets and tested on the 20%. (B) Average cross-prediction model AUC values for the different models when trained on the x-axis datasets. (C) Average AUC values for the different models considered when we leave one dataset out (x-axis) and train the model on the other three. We performed a 10-fold cross validation repeated 100 times and reported the average AUC values as shown. (D) Number of important features identified in the LODO settings using random forest when the model was trained on the other three and the country shown is exempted from training with their AUC values when predicted with those features for CAIM and CAIM_abun. (B and C). (E) Bump charts illustrated the top five important features in all the RF models trained for the LODO setting and their ranks in the other scenarios (y-axis). X-axis represents the training sets or datasets left out of the model. (F) Compariaionm of AUC of prediction models of LASSO, SVM and RF based on taxonomic profiles generated by CAIM and MetaPhlAn3 of the Thai primary liver cancer dataset.

Among the three predictive models, the predictive performance of the RF model in discriminating between CRC patients and healthy controls outperformed the other predictive models for both CAIM and CAIM_abun. Using CAIM as the input for RF model construction improved the predictive performance (AUC) over that of previous studies [21, 36, 37]. Thus, using RF in the LODO setting, we evaluated the predictive performance of highly important features as biomarkers for discriminating CRC from controls (Fig. 5D). When using the top 16 most important features, the predictive model was able to achieve a higher AUC value than when using all the features, except for when USA datasets were used. This poor performance on the USA dataset, compared to the other countries’ datasets, was also observed in the cross-prediction and the LODO setting. Next, we assessed the consistency of these predictive features as biomarkers for early screening or detection of CRC in patients using CAIM and CAIM_abun. Based on the top five important features from the RF model in the LODO setting (Fig. 5F), we considered their order of importance or rank in the models. The features identified with CAIM were more consistent across the different LODO predictive model scenarios than those identified by CAIM_abun. Most of the top five features identified using CAIM ranked within 1–7, while those that were identified with CAIM_abun ranked within 1–66. There was also some consistency in the order of importance for some of the species (*Parvimonas micra, Fusobacterium nucleatum,* and *Gemella morbillorum*) using CAIM across the datasets, but not with CAIM_abun. These results strongly indicated that using the genome-coverage cutoff provided a more reliable species biomarker. In addition, we peformed three classes predictive model using taxonomic profiling results obtained from either CAIM and MetaPhlAn3 [39], which was use in the original study, identified from WMS of stool amples from Thai primary liver cancer cohort [24] to predict the three classes of intrahepatic cholangiocarcinoma, hepatocellular carcinoma and healthy control. We found that predictive models using taxonomic profiling from CAIM gave a better performance (Fig. 5F). In this dataset SVM formulation give the best results compared to LASSO and RF.

## DISCUSSION

The use of genome coverage analysis for identification of microbiome (CAIM) improved taxonomic classification of metagenomic samples. The refinement step implemented in CAIM validates the reference genomes selected for alignment. CAIM is an alignment-based technique; thus, the refinement step helps reduce the number of reference genomes that are selected to be contained in the metagenomic reads, as there is no clear consensus on what a good containment-index cutoff should be, thus ensuring that only highly likely selected true reference genomes are used in the alignment. We recommend further research into the appropriate containment-index cutoff. Additionally, using the sequencing depth information for a microbial taxa in CAIM improved the accuracy of abundance estimates for the perfect scenario (Fig. 2C) and throughout the CAIM pipeline (Fig. 3B).

The use of the genome coverage in CAIM tremendously improved the F1 score compared to using the relative abundance (Fig. 2D). Most of the false-positive microbial taxa had a relative abundance passed the cutoff howere, they had a low percentage of its genome size covered by the reads in the mock community dataset. This was the key reason for CAIM_abun underperforming CAIM in terms of the average F1 score when we varied both the genome coverage and relative-abundance cutoff. For the staggered genomic mix mock community dataset, we observed a low genome coverage for species with low percentages of genomic DNA, resulting in a genome-coverage cutoff of 15% missing those species. CAIM also identified some viral species and phages (*Staphylococcus virus*, *Staphylococcus phage,* and *Pseudomonas phage*) with a high genome coverage in most of the bacterial mock community ATCC samples for both short and long reads. *Cellulophaga baltica* and *Pseudoalteromonas marina* were the most common bacteria identified by CAIM from the viral mock community, because the dsDNA viruses were hosted by either the *Cellulophaga baltica* or *Pseudoalteromonas,* while the ssDNA viruses were propagated *on Escherichia coli* [40].

Building predictive models using relative abundance values obtained with CAIM had a better performance compared with CAIM_abun. Filtering false positives with the genome coverage rather than the relative abundance improved the predictive power of the models in most of the scenarios considered. Thus, we recommend the use of a genome-coverage cutoff of 5% for real metagenomic datasets as we observed that it improves discriminating CRC patients from healthy controls compared to a genome-coverage cutoff of 15%. Evaluation of different genome-coverage cutoffs for different metagnome data is recommended. Further investigation is required to define a genome-coverage cutoff for complex environmental metagenomic samples. Most of the important features or microbial species (e.g., *Parvimonas micra, Fusobacterium nucleatum*, *Solobacterium moorei, Gemella morbillorum*) that we found have been established as known biomarkers for discriminating CRC patients from controls [21, 23, 37, 38, 41]. Using CAIM with genome-coverage filtering of false positives was more robust than using the relative-abundance cutoff with and without the refinement step.

## CONCLUSIONS

We propose a new alignment-based taxonomic classifier for identifying microbial taxa present in a metagenomic sample using the genome coverage as a pre-filter and the sequencing depth for estimating the relative abundance. We used two different containment techniques, with one serving as a validation of the results from the other to identify the true reference genomes in metagenomic samples. Using simulated and real datasets, we demonstrated the strength of using CAIM in identifying microbial taxa compared with other classifiers, CAIM consistently achieved high F1 scores across various samples.

## Software availability

CAIM is publicly available at GitLab (https://gitlab.com/Dacheamp/caim) under an MIT license.

The WMS data generated by this study is available at NCBI Sequence Read Archive database under bioproject PRJNA1000750

## Author contribution

I.N. designed and conceived the project. D.A developed and implemented the CAIM software. D.A., P.J., A.K., Y.P. and I.N. performed computational analysis. T.W., A.K.,S.K., P.K. and N.C. performed matgeneomic sequencings under the supervision of I.N. Y.P.,P.J., A.K. and K.K. perform rigorous testing and improvement on CAIM software. D.A. and I.N. wrote and edited the manuscript, and all authors edited the manuscript. All authors have read and approved the final version.

## Acknowledgement

We thank Mr. Pantakan Puengrang for his technical assistance in beginning of the study.

