## Supplementary Material for "CAIM: Coverage-based Analysis for Identification of Microbiome"

**Representative reference genomes database**

To construct a representative reference genome database, we downloaded prokaryotic and archaea genomes from the Genome Taxonomy Database (03/23/2022) [1], and fungi genomes were downloaded from the joint genome institute (JGI) (11/19/2021) [2] , as well as the National Center for Biotechnology Information (NCBI) RefSeq (09/12/2021) [3] for those not present in JGI. All complete NCBI RefSeq viral genomes [4] were also included in our reference database. To allow for a fair comparison with other tools, we included genomes for which there were no complete assemblies present in the NCBI genome database. Taxonkit [5] was used to assign taxonomy for all the reference genomes using their taxid. To reduce the large numbers of reference genomes, we kept all genomes with taxonomic information to the strain level. For multiple species without strain information, we selected species with the largest genome size as part of the representative reference genome database. There were 41,386 genomes in the representative reference database—1,346 archaea genomes, 27,658 bacteria genomes, 1,754 fungi genomes, and 10,628 viral genomes. We assembled a perfect representative reference database using representative genome of each species present in the mock community (MC) datasets to create a perfect scenario of the representative reference database for each mock dataset. A mock community in this study is a community of known taxonomic information and their respective genomic information used in constructing the community.

**In-house datasets: Sample collection**

Two types of samples were used: a mock community standard (Zymo Research, USA) and five stool samples. The mock microbial community, which was comprised of two fungi and eight bacteria, was used to validate a DNA isolation protocol and construct a Nanopore sequencing analysis workflow. The five stool samples used in this study were obtained from C57BL/6J mice (Jackson Laboratory, USA), athymic nude mice (Foxn1nu) (Jackson Laboratory, USA), a healthy female dog, and a healthy man and woman. All fresh stool samples (200 mg) were collected into sterile containers (Falcon™, USA) and transferred immediately to DNA/RNA Shield™ Lysis Tubes (Zymo Research, USA) using a sterile spoon (Corning, USA). All the lysis tubes were further homogenized using Disruptor Genie™ (Zymo Research, USA).

**Genomic DNA extraction**

The DNA was extracted from each sample using the ZymoBIOMICS™ DNA extraction protocol (Zymo Research, USA). The DNA purity was quantified using a Nanodrop Spectrophotometer (NanoDrop Technology, USA) by measuring the A260/280 and A260/230 ratios. The DNA concentration was quantified using a Qubit® 3.0 Fluorometer (Invitrogen, USA). The integrity of the extracted DNA was evaluated with agarose gel electrophoresis (Thermo Scientific, USA). Extracted DNA samples were stored at 4 °C for 24 hours before sequencing.

**Library preparation and genomic DNA sequencing by Oxford Nanopore Technology**

*Native metagenome seqeuncig :* We used a MinION (Oxford Nanopore Technologies [ONT], USA) device with a ligation sequencing kit (SQK-LSK108, ONT, USA) and followed the manufacturer's 1D Genomic DNA sequencing protocol for library preparation. Briefly, for each sample, 1100 ng of genomic DNA was repaired using the NEBNext FFPE repair mix (New England Biolabs [NEB], USA), according to manufacturer recommendations, and the DNA was subsequently purified using AMPureXP beads (Beckman Coulter, USA). End-repair of DNA fragments was then performed using the NEBNext Ultra II End Prep/dA-tailing module (NEB, USA), and DNA was subsequently purified again using AMPure XP beads. Next, ligation of motor protein-complexed AMX adapter to genomic DNA ends was carried out using the NEB

Blunt/TA ligase Master Mix (NEB, USA). A final round of AMPure XP purification was then performed before the DNA library was eluted and loaded onto a flow cell for sequencing. Sequencing of the genomic DNA was performed on a single R9.5/FLO-MIN107 flow cell (ONT, USA) on a MinION Mk1B (ONT, USA) for 48 hours.

*Metagenome amplification and seqeuncig* : A total of 220−600 ng of genomic DNA, extracted from human fecal samples, was performed rolling cycle amplification (RCA) using phi29-XT DNA polymerase and exonuclease-resistant random primers (New England Biolabs, MA, USA). The reactions were incubated at 42 °C for 4 h, followed by heat inactivated at 65 °C for 10 min. The resulting RCA products were purified using 1X AMPure XP beads (Beckman Coulter, CA, USA), and 3 µg of the purified RCA was subsequently debranched using 30 units of T7 Endonuclease I (New England Biolabs, MA, USA). The reactions were then incubated at 37 °C for 2h. After 1X AMPure XP beads purification, the debranched DNA were carried out library preparation for nanopore sequencing using Native Barcoding Kit SQK-NBD114.24 (Oxford Nanopore, Oxford, UK), and sequenced using a MinION and R10.4.1 flow cells (Oxford Nanopore, Oxford, UK).

Base-calling was performed using the either Guppy version 4.6.0 local-based software (ONT, USA). All FASTQ data were trimmed with Porechop version 0.2.4 (https://github.com/rrwick/Porechop) to remove adapters from read ends and split sequences with internal adapters. Nanofilt (https://github.com/wdecoster/nanofilt) version 2.3.0 was used to filter out reads shorter than 200 bp and with a mean quality of less than 8.

**Library preparation and genomic DNA sequencing by Illumina MiSeq**

Each extracted genomic DNA sample was adjusted to 0.2 ng/µL and then prepared for sequencing with a Nextera XT DNA Sample Prep Kit (Illumina Inc., USA). In total, 1 ng of each sample was used for tagmentation, as directed by the manufacturer. After tagmentation, PCR amplification was done using a unique combination of barcode primers for each of the 6 samples. Then, each DNA library was purified using AMPure XP beads (Beckman Coulter, USA) and normalized with Library Normalization beads/additives. Normalized libraries were pooled, and pair-end sequencing was performed using the MiSeq Reagent Kit v3 (300 cycles) through the Illumina MiSeq platform.

**Public mock community datasets**

Several publicly available and in-house mock community datasets were used in this research. There were 30 mock community datasets in total assembled with Illumina, PacBio, and Nanopore sequencing platforms (Supplementary TableS1). The sizes of the mock community datasets ranged from 1.2 Gb to 136 Gb (Fig 2A). The mock community datasets consisted of 18 short-read datasets sequenced on the Illumina platform and 12 long-read datasets sequenced on both Nanopore and PacBio platforms. The viral mock community MCA-G2, MCA-N1, MCA-S1, MCB-G1, MCB-N1, and MCB-S1 datasets were composed of 10 dsDNA viruses and 2 ssDNA viruses but with different compositions sequenced using three different Illumina MiSeq platform protocols ("G" for Mulitple displacement amplification (MDA), "N" for Tagmentation (TAG), and "S" for Adaptase-Linker amplification (A-LA)). Detailed information on how the community was generated has been published elsewhere [6]. The MCA mocks had a low total abundance of ssDNA viruses (~2% of the community), while the MCB mocks had a high total abundance of ssDNA viruses (~66% of the community). The abundances of the species composition for the viral mocks ranged between 0.15% - 58.62%, 0% - 100%, 0.22% - 26.12%, 0.01% - 58.07%, 0.07% - 26.33%, and 0.24% -58.18% for MCA-G2, MCA-N1, MCA-S1, MCB-G1, MCB-N1, and MCB-S1, respectively.

The Illumina metagenomic mock datasets for Gut-HiLo, Gut-Mix, ATCC MSA-1000, ATCC MSA-1001, ATCC MSA-1002, and ATCC MSA-1003 were obtained from the NCBI under Bioproject ID PRJNA622674, details of which can be found elsewhere [7] (Supplementary Table 1). The datasets large consisted of bacterial species. The Gut-HiLo mock communities were staggered composition of 20 gut microbiome strains with genomic dna composition ranging from 0.11% - 19.82% and the Gut-Mix mock communities were even compositions of 20 gut microbiome strains with genomic DNA composition ranging from 0.11% - 19.82% and 2.69% - 8.97%. The two *Bifidobacterium longum* strains (infantis and longum) were considered one species whose composition was the sum of two strains' compositions, thus reducing the species to 19 for both the Gut-HiLo and Gut-Mix. The ATCC MSA-1000 mock community dataset consisted of 10 species with an even genomic DNA composition of 10%. The ATCC MSA-1001 mock community dataset consisted of a 10-strain staggered mix of genomic DNA ranging from 0.04% - 44.78%. There were two ATCC MSA-1002 mock community datasets sequenced on the Illumina platform (PRJNA622674), and our in-house data was sequenced on the Nanopore sequencing platform. The ATCC MSA-1002 mock consisted of a 5% even strain genomic mix of 20 species. There were 3 ATCC MSA-1003 mock community datasets sequenced on Illumina, Nanopore, and PacBio platforms. The PacBio ATCC MSA-1003 [8] mock contained HiFi reads generated using the Sequel II System. The ATCC MSA-1003 mock consisted of a 20-strain staggered mix of genomic DNA with a composition ranging between 0.02% - 18%.

Several ZymoBIOMICS (Zymo) mock community datasets across different sequencing platforms were used in this work. The ZymoBIOMICS D6300 Microbial Community Standard samples used in this study (ZymoIL, Zymo-R103, ZymoQ20, Zymo-LOG, and Zymo-EVEN) all had the same 10 species and abundance proportions. The ZymoIL sample was sequenced on the

Illumina platform and the others on the Nanopore platform. The abundance composition for the species in these samples ranged between 2% - 12%. Zymo-R103, Zymo-LOG, and Zymo-EVEN were generated on the Oxford Nanopore GRIDION, while ZymoQ20 was generated on the Oxford Nanopore ProMETHION with Q20 chemistry. ZymoBIOMICS D6331 Gut Microbial Community Standard labeled (ZymoD6331) is a PacBio HiFi dataset, which contains 17 species with abundance composition ranging from 0.0001% - 14%. The three different strains of *Escherichia coli* were considered one species with their abundance composition summed. Any genus *Shigella* species were also counted towards *Escherichia coli* due to their high genomic sequence similarities. For ZymoBIOMICS mock communities, we filtered out all the viral species detected by the taxonomic classifiers. For samples with Bacillus subtilis, most of the tools identified Bacillus intestinalis at high abundance, thus the analysis was done at the genus Bacillus.

We also obtained BBMock-12, a publicly available 12-bacteria strain mock community and a 20 species mock community (GIS20), both sequenced on the Illumina, Oxford Nanopore, and PacBio platforms. Detailed information about BBMock-12 and GIS20 has been previously published [9] respectively. The staggered abundance of the 12 species varies from 0.9% - 16.4%. There were no genome assemblies for *Thioclava sp. ES.032*, *Psychrobacter sp. LV10R520-6,* or *Marinobacter sp. LV10R510*-8; thus, we analyzed this dataset at the genus level. The abundance of the 20 species for the GIS20 ranges from 0.1 – 30%. FungiMock [10], a fungal mock community dataset sequenced with the Oxford Nanopore and Illumina platforms, was obtained and used in this study and labeled as Mock-Fungal. This dataset consisted of 43 species but 2 (*Diutina mesorugosa* and *Cryptococcus magnus*) did not have genome assembly; thus, the analysis was done at the genus level with 24 genera.

**Synthetic and real datasets**

We downloaded synthetic data used in the Critical Assessment of Metagenome Interpretation (CAMI) [11] competition to assess the performance of our classifier. We used the low and medium complexity (sample 1 270bp insert) of their simulated datasets. The real datasets were generated in our lab on both the Oxford Nanopore and Illumina sequencing, as described above. Datasets sequenced in our lab are available at NCBI with project id PRJNA1000750.

**Genomic coverage and relative abundance estimation**

Sequencing depth and genomic coverage play a crucial role in estimating the relative abundance and identification of organisms in the metagenomic sample [12]. We estimated genomic coverage ($C$) as the proportion of the genome size covered by the aligned reads expressed in percentage [12]. Thus, for a given genome of size ($l$) with $m$ length covered by the aligned reads, $C = 100 * m/l$. To estimate the empirical optimal genome coverage cutoff for filtering out false positives in our classifier, we subsampled 50 metagenomic samples from the various mock community datasets for both short and long reads with varying sequencing depths (629M – 5G) using SeqKit [13], except for the datasets that were missing some genome assembly in the reference database. Varying the genome coverages for each subsampled mock dataset, we estimated their F1 score, and the coverage cutoff that ensured a balance between the precision and recall in both the short and long reads was used as the default cutoff value. To estimate the relative abundance of genomes in a sample, we followed previously published approaches [47, 48] but with varying sequencing depths. We defined the sequencing depth of a reference genome to be the number of nucleotides that aligned to the reference genome divided by the genome size. For a given mixture of sequenced metagenomic samples, we define the total nucleotides $T = \sum_i^m T_i,\ i = 1,2,\ldots,m$ , where $T_i$ are the total nucleotides that aligned to the genome $i$ and $m$ are the total number of organisms present in the sample. For genome $i$, $T_i = \beta_i l_i$ where $\beta_i$ is sequencing depth for genome $i$. The sequencing depth of a genome is proportional to the relative abundance of the genome. We define the relative abundance (nucleotide abundance) of a genome $i$ as $a_i = \frac{\beta_i}{\sum_i^m \beta_i}$ . We compared our nucleotide-abundance approach of estimating the relative abundance to the traditional read-count approach—the number of reads that mapped to a genome divided by the total sequencing reads.

**Taxonomic classification method comparison**

Four short-read taxonomic classifiers and two long-read taxonomic classifiers were used in this study for benchmarking. Several works have benchmarked the performance of most of these classifiers [8]. We could not include some tools in this work as we found them to be obsolete, no longer maintained by the authors, or inoperable. For long-read metagenomic samples, we compared CAIM with Kraken2. For short-read metagenomic samples, we compared CAIM to Metalign, StrainPro, KMCP, and Kraken2. Metalign is an alignment-based approach that maps short reads to a pre-filter nucleotide reference database obtained by the min hash approach. KMCP is a k–mer-based approach for profiling metagenomic reads against a chunked reference database. Kraken2 is a k–mer-based approach, whereas StrainPro is an alignment-based approach for assigning microbial taxa to metagenomic samples.

**Evaluation**

We evaluated the performance of our method in comparison with other existing methods on synthetic and several long- and short-read datasets on the presence/absence of organisms at the taxonomic levels and the relative abundance estimation. Recall, precision, and F1 score were the detection metrics used to assess the performance of the methods. Recall is the proportion of true positive organisms that were actually predicted to be present in the sample, and precision is the proportion of predicted organisms that are actual true positives in the sample [14]. The numerical error measure, root mean square error (RMSE), [15] was used to assess the accuracy of the relative abundance estimation approaches. We also evaluated the performance of these tools to identify species at various abundance cutoffs as compared to our approach using a fixed genome coverage value.

Area under the receiver operating characteristic curve (AUC) score values were used to evaluate the performance of the prediction models. The AUC has been used extensively in evaluating prediction models [16]. A high AUC value implies the model good at discriminating between species. Random forest (RF), support vector machine (SVM), and least absolute shrinkage and selection operator (LASSO) were the three models evaluated on the CRC and healthy control fecal metagenomic samples in this study.

**Fecal metagenomic colorectal cancer datasets and biomarker analysis**

Publicly available fecal shotgun metagenomic colorectal (CRC) and healthy datasets from 4 different countries were downloaded and used in this study. For detailed information on the datasets and the criteria for the inclusion and exclusion of a sample, kindly refer to Thomas et al. [17]. All fastq files were downloaded from the European Nucleotide Archive (ENA) with the identifiers PRJEB10878 for CHINA [18], ERP008729 for AUSTRIA [19], PRJEB12449 for USA [20] and ERP005534 for GERMANY [21] The Austria, China, Germany, and USA fecal metagenomic datasets consisted of 46, 74, 60, and 52 CRC patients and 63, 54, 60, and 52 healthy patients, respectively (Supp. Table 3). Species identification and relative abundance estimation for the CRC patients and healthy controls were done with CAIM and CAIM_abun. For CAIM_abun, we filtered out all species with relative abundances less than 0.01, and a coverage cutoff of 5 was used for CAIM. Gao et al. [22] used similar datasets and found that correcting for batch effects before training predictive models did not improve their predictive performance; thus, we also decided not to control for batch effects.

We built and trained predictive models on the relative-abundance profiles to investigate if species identified with CAIM and CAIM_abun could help discriminate between CRC patients and healthy controls and which method had a better performance. For within and cross-validations, we filtered out species with the number of nonzero abundances less than 1% of the sample size for each dataset. The relative-abundance profile table was log10 transformed after adding a small pseudo count of 1e-10 and later standardized as z-scores. Using the standardized z-scores, we split the dataset into 80-20, where we train our models on 80% of the dataset for 100 repeats of 10-fold cross-validation and predict the remaining 20% for the within-sample validation. We compared the performance of the models using AUC for CAIM and CAIM_abun. For each prediction, we estimated and averaged the AUC values for the 30 repetitions. RF, SVM, and LASSO models were built and used for the prediction. The RF model used 1,000 decision trees with a random search.

In addition were perform three clsses analysis of our stool samples of liver cancer cohorts [23] of intrahepatic cholangiocarcinoma (iCCA), hepatocellular carcinoma (HCC) and healthy control. We compare the AUC of prediction models of LASSO, SVM and RF based on taxonomic identification and profiles generated by CAIM and MetaPhlAn3 [24]. For each prediction, we estimated and averaged the AUC values for the 100 repetitions. RF, SVM, and LASSO models were built and used for the prediction. The RF model used 1,000 decision trees with a random search.

**Comparative analysis of taxonomic difentifiction between Nanopor and Illumnia sequencing platform using stool samples of different host species using CAIM.**

The sequence datasets of generated by Nanopore and llumina Platform obtained from the sample aliquot were analyzed by CAIM to genate taxonomic profiles. The profiles were import to R and phyloseq package [25] for futhur comparisons. The correlation analysis of species relative abundance btween the two platforms were peformed using spearman rank correlation. The alpha diversing analysis using Shannon and Simpson diversity index were calculated for indivaul sample. The beta diversing analysis using Non-metric multidimensional scaling (NMDS) based on Bray-Cutis distance were peformed across the samples and the result were ploted in two demnsional with eclipse statistic of individual host species.

**Data Filtering and Analysis**

All long reads with read lengths less than 200 base pairs and quality less than 8 were filtered out using NanoFilt [26]. For short Illumina reads, we used BBduk [27] with the following parameters *ktrim=r, k=21, mink=10, tpe tbo, qtrim=r, trimq=20 and minlen=30* to filter out poor quality short reads. For taxonomic classification of metagenomic samples, we used the default parameters and in-built database for each taxonomic tool with the exception of Kraken2 and StrainPro for which we built a reference database using the representative reference genomes used in CAIM. All the predictive models were built in R (version 4.1.0) using the CARET package [28].
